# Genomic evidence that governmentally produced *Cannabis sativa* poorly represents genetic variation available in state markets

**DOI:** 10.1101/2021.02.13.431041

**Authors:** Daniela Vergara, Ezra L. Huscher, Kyle G. Keepers, Rahul Pisupati, Anna L. Schwabe, Mitchell E. McGlaughlin, Nolan C. Kane

**Affiliations:** Kane Laboratory, Department of Ecology and Evolutionary Biology, University of Colorado Boulder, Boulder, Colorado, USA; Gregor Mendel Institute (GMI), Austrian Academy of Sciences, Vienna Biocenter (VBC), Dr. Bohr-Gasse 3, 1030 Vienna, Austria; University of Northern Colorado, School of Biological Sciences, Greeley, CO 80639, USA

**Keywords:** cannabinoids, copy number variation, genome diversity, hemp, repetitive genomic content, marijuana, NIDA, THC

## Abstract

The National Institute on Drug Abuse (NIDA) is the sole producer of *Cannabis* for research purposes in the United States, including medical investigation. Previous research established that cannabinoid profiles in the NIDA varieties lacked diversity and potency relative to the *Cannabis* produced commercially. Additionally, microsatellite marker analyses have established that the NIDA varieties are genetically divergent form varieties produced in the private legal market. Here, we analyzed the genome of multiple *Cannabis* varieties from diverse lineages including two produced by NIDA, and we provide further support that NIDA’s varieties differ from widely available medical, recreational, or industrial *Cannabis*. Furthermore, our results suggest that NIDA’s varieties lack diversity in the single copy portion of the genome, the maternally inherited genomes, the cannabinoid genes, and in the repetitive content of the genome. Therefore, results based on NIDA’s varieties are not generalizable regarding the effects of *Cannabis* after consumption. For medical research to be relevant, material that is more widely used would have to be studied. Clearly, having research to date dominated by a single, non-representative source of *Cannabis* has hindered scientific investigation.

## Introduction

Public perception of recreational and medicinal *Cannabis* use has shifted, with *Cannabis* derived products quickly becoming a multibillion-dollar legal industry. However, the National Institute on Drug Abuse (NIDA), a United States (U.S.) governmental agency, continues to be the sole producer of *Cannabis* for research. Additionally, high-THC producing *Cannabis* continues to be classified as a Schedule I drug, along with heroin, LSD, and ecstasy, according to the DEA (DEA 2020). This schedule I classification restricts the acquisition of *Cannabis* from the private markets, making NIDA the only federally legal source for research. In addition to this limitation, research on *Cannabis* requires a multitude of permits and supervision (Nutt et al. 2013; Hutchison et al. 2019). However, the medical and recreational *Cannabis* industry in North America are predicted to grow to 7.7 and 14.9 billion dollars, respectively, by late 2021 (Hutchison et al. 2019).

*Cannabis sativa* L. (marijuana, hemp) is an angiosperm member of the family Cannabaceae (Bell et al. 2010). It appears to be one of the oldest domesticated plants, utilized by numerous ancient cultures, including Egyptians, Chinese, Greeks, and Romans (Li 1973, 1974; Russo 2007). This versatile plant has many known uses, including fiber for paper, rope and clothing, oil for cooking and consumption, and numerous medicinal applications. The plant produces secondary metabolites known as cannabinoids that interact with the human body in physiological (Russo 2011; Swift et al. 2013; Volkow et al. 2014) and psychoactive (Russo and McPartland 2003; ElSohly and Slade 2005) ways. Cannabinoids are terpenoid compounds (Zwenger and Basu 2008) that are concentrated in the trichomes of the female flowers (Sirikantaramas et al. 2005). The remarkable properties of cannabinoids are partly responsible for driving the growth of the thriving *Cannabis* industry. Two of the main cannabinoids--Δ-9-tetrahydrocannabinolic acid (THCA) and cannabidiolic acid (CBDA)--when heated are converted to the neutral forms Δ-9 tetrahydrocannabinol (THC) and cannabidiol (CBD), respectively (Russo 2011). The most well-characterized enzymes responsible for the production of these cannabinoids in the plant are Δ-9-tetrahydrocannabinolic acid synthase (THCAS) and cannabidiolic acid synthase (CBDAS).

Despite the regulatory hurdles and the limited scope of contributions from the U.S. government, *Cannabis* research is growing at a rapid pace (Vergara et al. 2016; Kovalchuk et al. 2020) and U.S. scientists have made significant advances in *Cannabis* research from multiple disciplines. Researchers in the U.S. have produced one of the most complete publicly available *Cannabis* genome assemblies to date, along with the locations of the cannabinoid family of genes in the genome (Grassa et al. 2018). However, oversight is needed to assure the quality and consistency of *Cannabis* testing across laboratories (Jikomes and Zoorob 2018). Regulation and supervision will allow for a deeper understanding about all of the compounds produced by the plant particularly minor cannabinoids which are not always measured (Vergara et al. 2020) and that have multiple genes related to their production with complex interactions (Vergara et al. 2019). This is particularly important because medical *Cannabis* use has outpaced its research (Hutchison et al. 2019). Collaborative research between American academic institutions and private companies has shown that the cannabinoid content and genetic profile of *Cannabis* provided by NIDA is not reflective of what consumers have access to from the private markets, therefore research with these varieties is discordant (Vergara et al. 2017; Schwabe et al. 2019).

In 2017, we compared the cannabinoid chemotypes from the *Cannabis* produced in the private market to the chemotypes from the governmentally produced *Cannabis* for NIDA by the University of Mississippi (Vergara et al. 2017). We found that NIDA’s *Cannabis* lacked potency and chemotypic variation and had an excess of cannabinol (CBN), which is a degradation product of THC. The cannabinoid diversity from the governmentally produced *Cannabis* was a fraction of that from the private markets. A study using microsatellite markers also showed that NIDA’s *Cannabis* was genetically different from commercially available recreational and medical varieties. This study concluded that results from research using flower material supplied by NIDA may not be comparable to consumer experiences with *Cannabis* from legal private markets (Schwabe et al. 2019).

Here, we present results of analysis to further examine the genetic diversity in governmentally produced *Cannabis*. We acquired DNA from two NIDA-produced samples which had been previously analyzed using ten variable microsatellite regions (Schwabe et al. 2019). After sequencing, we compared their overall genomic diversity to that of previously sequenced varieties including hemp and marijuana-types (Lynch et al. 2016; Vergara et al. 2019). We report here the genomic characteristics of the two NIDA samples, including overall genetic variation, as well as genetic variation within the cannabinoid family of genes, the maternally inherited organellar genomes (mitochondrial and chloroplast), and the repetitive genomic content. We compare this diversity to the publicly available genomes from other *Cannabis* lineages within the species, to characterize the relationships with other well-studied lineages.

## Materials and Methods

### NIDA’s samples

Bulk *Cannabis* supplied for research purposes is referred to as “research grade marijuana” by NIDA and is characterized by the level of THC and CBD (NIDA 2016). They offer 12 different categories of *Cannabis* for research that vary in the levels of THC (low < 1%, medium 1-5 %, high 5-10 %, very high >10%) and CBD (low < 1%, medium 1-5%, high 5-10%, very high > 10%)”. The high THC NIDA sample (Table S1) has an RTI log number 13494-22, reference number SAF 027355 and the high THC/CBD has an RTI log number 13784-1114-18-6, reference number SAF 027355. DNA from both samples was extracted by (Schwabe et al. 2019) and provided to the University of Colorado Boulder. These two samples were sequenced using standard Illumina multiplexed library preparation protocols as described in (Lynch et al. 2016) which yielded to an approximate coverage of 17-20x (Table S1).

### Genome assembly, whole genome libraries, and nuclear genome exploration

We aligned sequences from 73 different *C. sativa* plants to the previously developed CBDRx assembly Cs10 (Grassa et al. 2018). These genomes were sequenced using the Illumina platform by different groups (Table S1) and are, or will be, publicly available on GenBank. For detailed information on sequencing and the library preparation of the 57 genomes sequenced by our group at the University of Colorado Boulder please refer to Lynch et al., 2016. The remaining 16 genomes were sequenced and provided by different groups (Table S1), however most of these genomes have been previously used in other studies (Lynch et al. 2016; Vergara et al. 2019).

We aligned the 73 libraries to the CBDRx assembly using Burrows-Wheeler alignment (ver. 0.7.10-r789; Li and Durbin 2009), then calculated the depth of coverage using *samtools* (ver. 1.3.1-36-g613501f; Li et al. 2009) as described in Vergara et al. (2019). We used GATK (ver. 3.0) to determine single nucleotide polymorphisms (SNPs). We filtered for SNPs lying in the single-copy portion of the genome (Lynch et al. 2016) which resulted in 7,738,766 high-quality SNPs. The single-copy portion of the genome does not include repetitive sequences such as transposable elements or microsatellites. Subsequently, we were then able to estimate the expected coverage at single-copy sites as in Vergara et al. (2019). We performed a STRUCTURE analysis (ver. 2.3.4; Pritchard et al. 2000) with K=3 in accordance with previous research (Sawler et al. 2015; Lynch et al. 2016). With these STRUCTURE results, we then classified the different varieties into four different groupings: Broad-leaf marijuana type (BLMT), Narrow-leaf marijuana-type (NLMT), Hemp, and Hybrid (Table S2). Hybrid individuals had less than 60% population assignment probability to a particular population. We found 12 individuals in the BLMT group, 16 in the Hemp group, 14 in the Hybrid group, and 31 in the NLMT group. We then used SplitsTree (ver. SplitsTree4; Huson 1998) to visualize the relationships between the 73 individuals, VCFtools (ver. 4.0; Danecek et al. 2011) to calculate genome wide heterozygosity as measures of overall variation, and PLINK (ver. 1.07; Purcell et al. 2007) for a principal component analysis (PCA).

### Cannabinoid gene exploration

Using BLAST, we found 12 hits for putative CBDA/THCA synthase genes in the CBDRx assembly (Table S3) with more than 80% identity and an alignment length of greater than 1000bp. For this BLAST analysis, we used the CBCA synthase (Page and Stout 2017), the THCA synthase with accession number KP970852.1, and the CBDA synthase with accession number AB292682.1.

We estimated the gene copy-number (CN) for the cannabinoid genes (Vergara et al. 2019) and calculated summary statistics of the CN for each of the 12 genes by variety (Table S1). Differences in the estimated gene CN between the cultivars for each of the 12 cannabinoid synthases gene family were determined using one-way ANOVAs on the CN of each gene as a function of the lineages (BLMT, Hemp, Hybrid, NLMT), with a later *post hoc* analysis to establish one-to-one group differences using the R statistical platform (Team 2013).

### Maternally inherited genomes

We used the publicly available chloroplast (Vergara et al. 2015) and mitochondrial (White et al. 2016) genome assemblies to construct haplotype networks using PopART (ver. 1.7; Leigh and Bryant 2015) using only variants with a high quality score in the variant call file (VCF). The chloroplast and mitochondrial haplotype networks comprised 508 and 1,929 SNPs, respectively.

### Repetitive genomic content

We used Repeat Explorer (ver.2; Novak et al. 2010)to determine the repeat content in 71 of the 73 genomes (Pisupati et al. 2018). We decided to exclude ‘Jamaican Lion’ (NLMT) and ‘Feral Nebraska’ (hemp) genomes due to low-quality reads that led to dubious results. We estimated the repetitive content of the genome and annotating repeat families using custom python scripts (https://github.com/rbpisupati/nf-repeatexplorer.git).

## Results

### Nuclear genome exploration

Our analysis of the nuclear genome used 7,738,766 high-quality SNPs from the inferred single-copy portion of the genome. The STRUCTURE analysis (Figure 1, top panel) shows the population assignment probabilities for all 73 different varieties including both of NIDA’s varieties. This analysis established that NIDA’s samples cluster with both the hemp and NLMT groupings, with less than 60% in either group and therefore we categorized them as ‘hybrid’ (Table S2). This classification led to 12, 16, 14, and 31 individuals from the BLMT, Hemp, Hybrid, and NLMT groups, respectively. In other words, the 12 individuals that are part of the Hemp group had a population assignment probability of more than 60% to this group, as well as those assigned to the NLMT or BLMT groupings. However, those individuals with a probability of less than 60% to a particular population were assigned to the ‘hybrid’ category, which includes both of NIDAs samples. We color-code the hemp individuals in orange, the NLMT in blue, BLMT in purple, and the hybrid individuals in gray.

**Figure 1.**
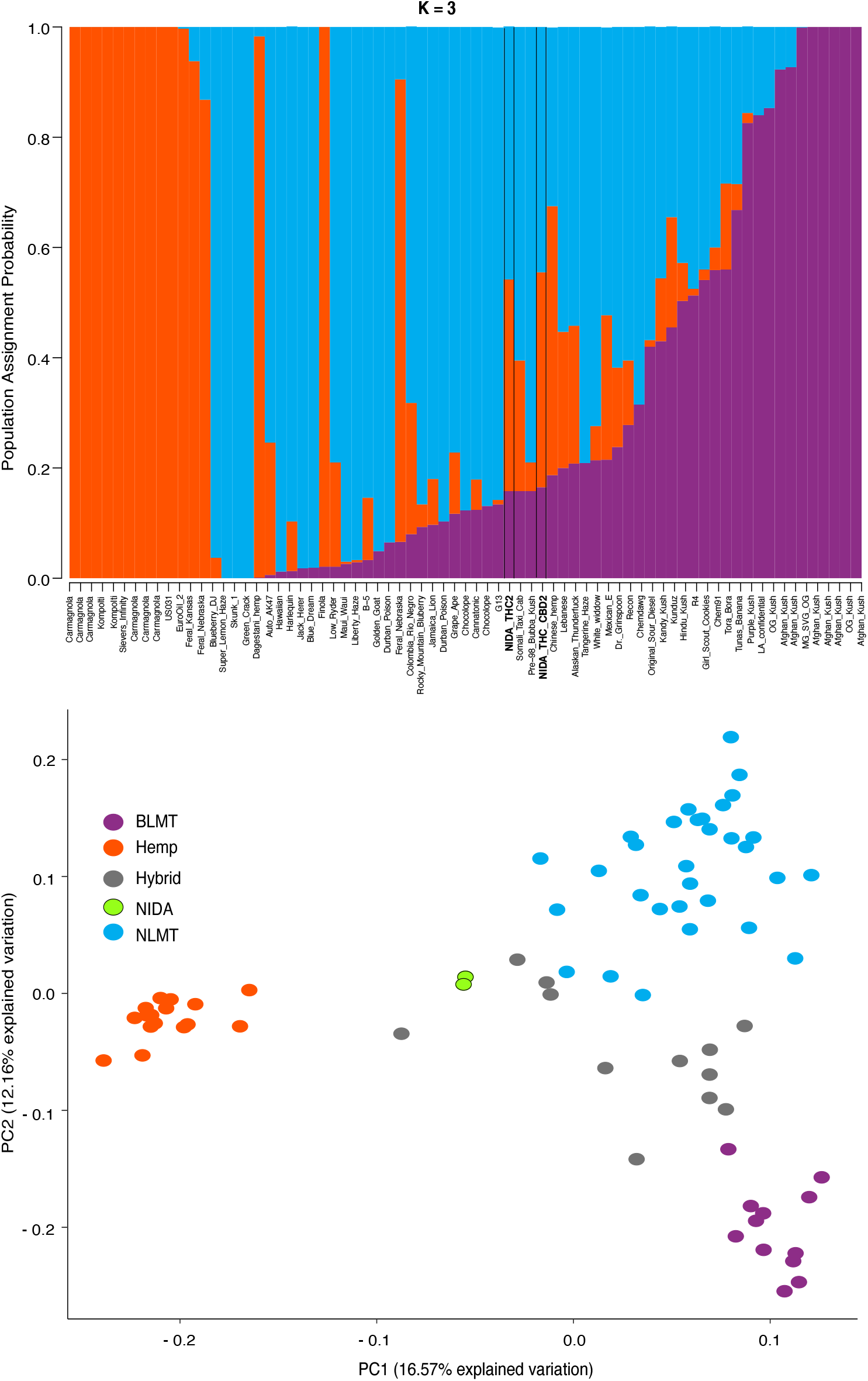
STRUCTURE and Principal Component Analyses. **Top panel:** Proportion of each color in the bar indicates the probability of assignment to Hemp (orange), NLMT (blue), or BLMT (purple), groups. Both of NIDA’s strains outlined with black margins are assigned to both NLMT and hemp groups with less than 60% probability, and therefore we assigned them to the Hybrid group. **Bottom panel:** The two NIDA varieties in green cluster with each other and away from other varieties.

In addition to clustering probability results (Figure 1 top panel) from STRUCTURE, we colored the varieties in the PCA (Figure 1 bottom panel) and SplitsTree (Figure 2) according to their population assignment probability. The first two principal components in the PCA explain 28.71% of the variation (Figure 1 bottom panel), and the two NIDA varieties cluster together, also seen in the SplitsTree analysis (Figure 2). Both the PCA and SplitsTree indicate high genetic similarity between the NIDA strains, and neither of them cluster with any other strains.

**Figure 2.**
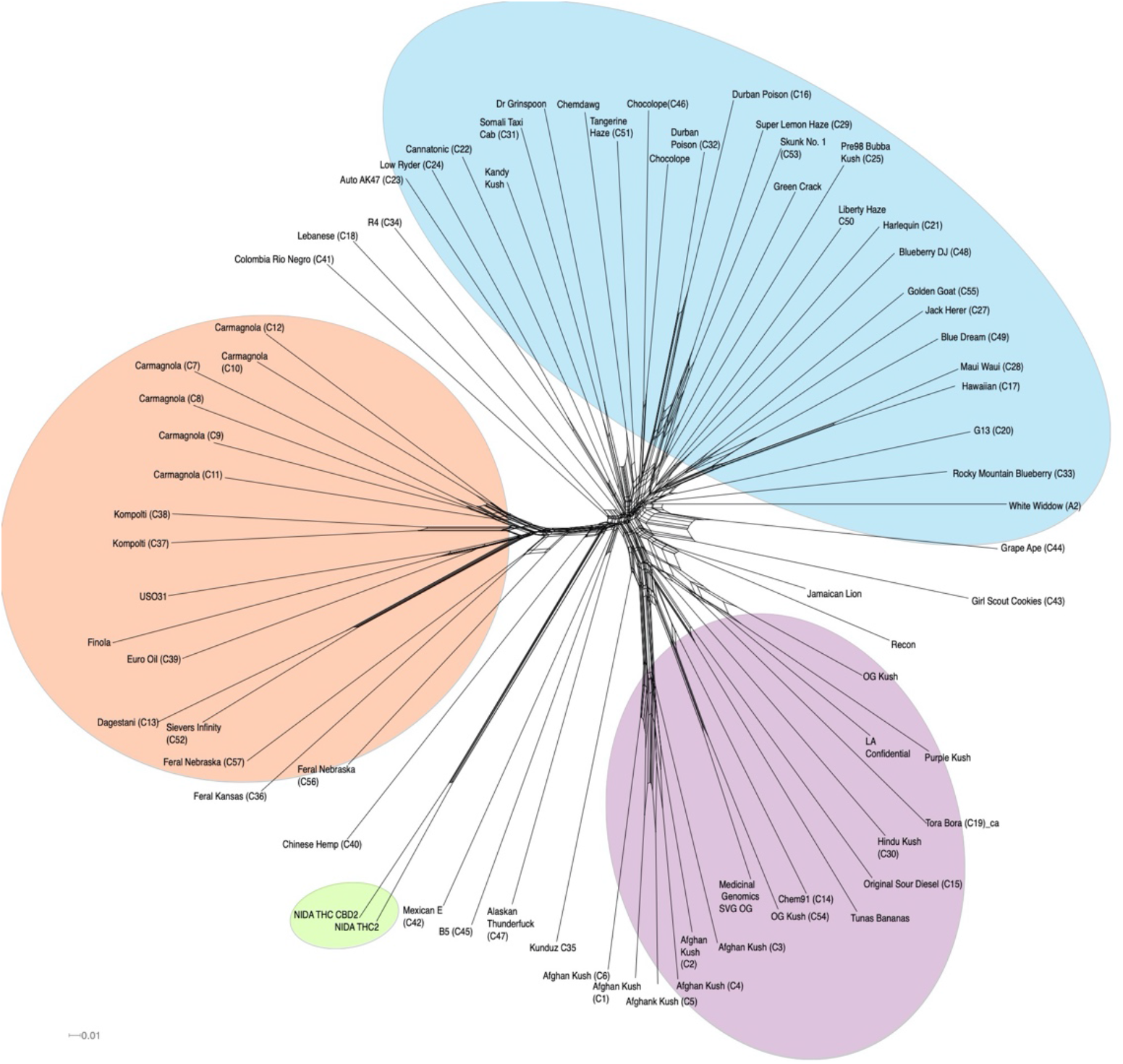
SplitsTree graph. Genetically similar individuals cluster together, such as the NIDA cluster, Afghan Kush cluster, and Carmagnola cluster. NIDA’s varieties highlighted in green. Hemp, NLMT, and BLMT shown in orange, blue, and purple respectively.

The hybrid group which contains NIDA’s varieties shows the widest range of heterozygosity (μ= 0.131; s.d= 0.0545) in the single-copy portion of the genome. However, it is not significantly different from any other group (Figure 3A). This wide range of heterozygosity in the hybrid group is expected given that we are grouping individuals that do not belong one particular genetic group but rather have some assignment probability to two or three genetic groups. Therefore, varieties which are not related to each other, or that belong to more than one group are found in the hybrid category. This is probably the reason why it is the highest of all other groups (hemp: μ=0.0817; s.d=0.0352; BLMT μ=0.0959; s.d=0.0405; NLMT μ=0.112; s.d=0.0411).

**Figure 3.**
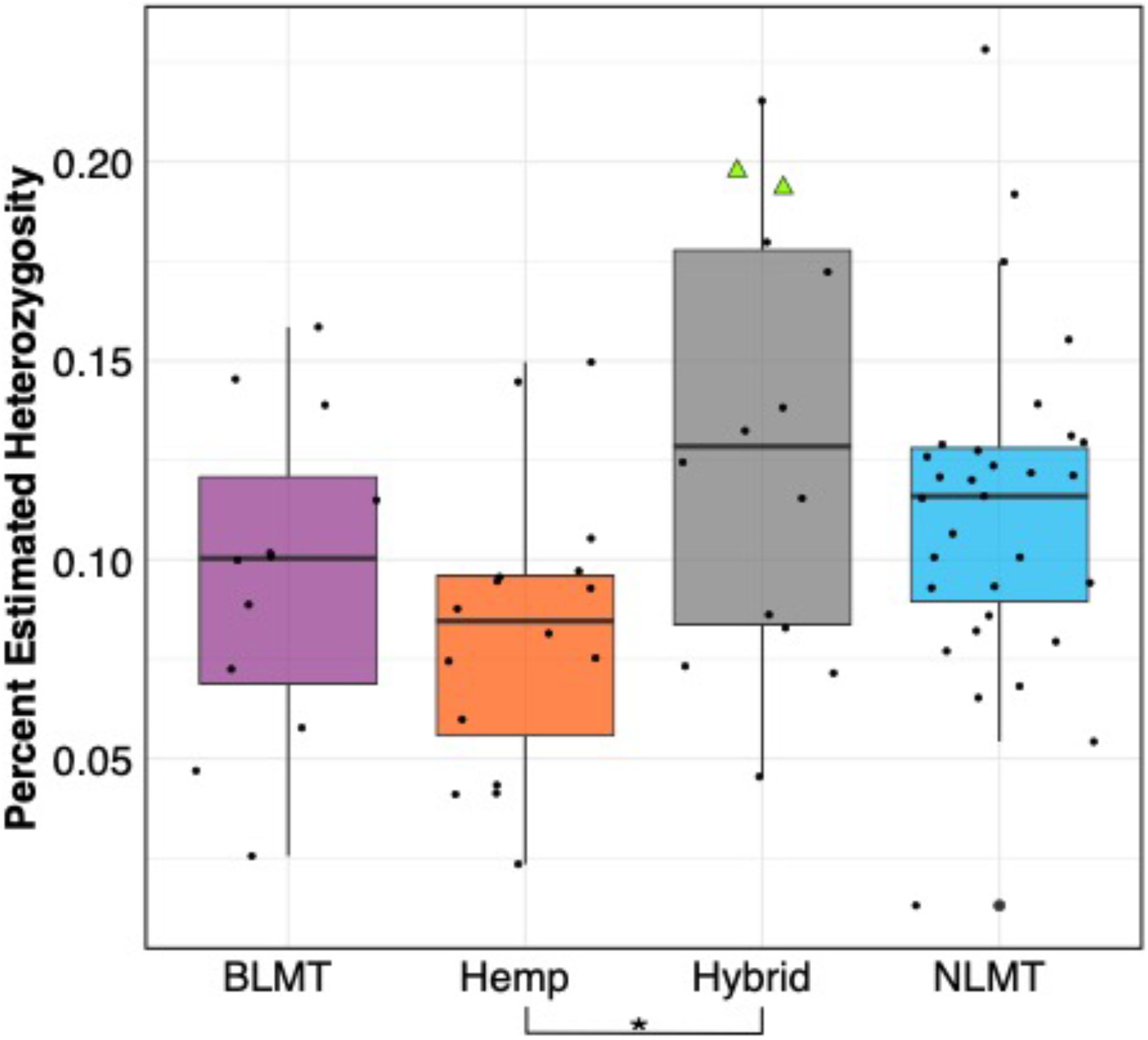
Genome wide heterozygosity. The hemp lineage differs significantly from the Hybrid grouping with a P<0.03. NIDA’s two varieties are presented within the hybrid grouping by two green triangles

### Cannabinoid gene exploration

Independent of which synthase we used for the BLAST analysis (either THCA, CBDA, or CBCA), the BLAST results delivered the same hits on the CBDRx assembly with different percent identities. Based on percent-identity scores, our BLAST results identified a hit in the CBDRx assembly that appears to code for cannabichromenic acid synthase (CBCAS), and one that possibly codes for CBDAS, but we did not find a hit for THCAS (Table S3). After calculating the copy number variation, we found that most groups have one copy of the CBCAS gene (BLMT μ=1.38; s.d=1.1; Hemp μ=1.88; s.d=2.15; Hybrid μ=1.56; s.d=1.33 and NLMT μ=1.44; s.d=2.57). Despite the hemp group having the widest range, no group significantly differed from each other (Figure 4A). For the CBCAS genes, the first NIDA sample has an estimated copy number of 0.37 and the second variety of 0.34. These values are on the lower side of the copy number distribution. We include the copy number variation of an unknown cannabinoid, which was the only other locus that had significant differences between groups (Figure 4B).

**Figure 4.**
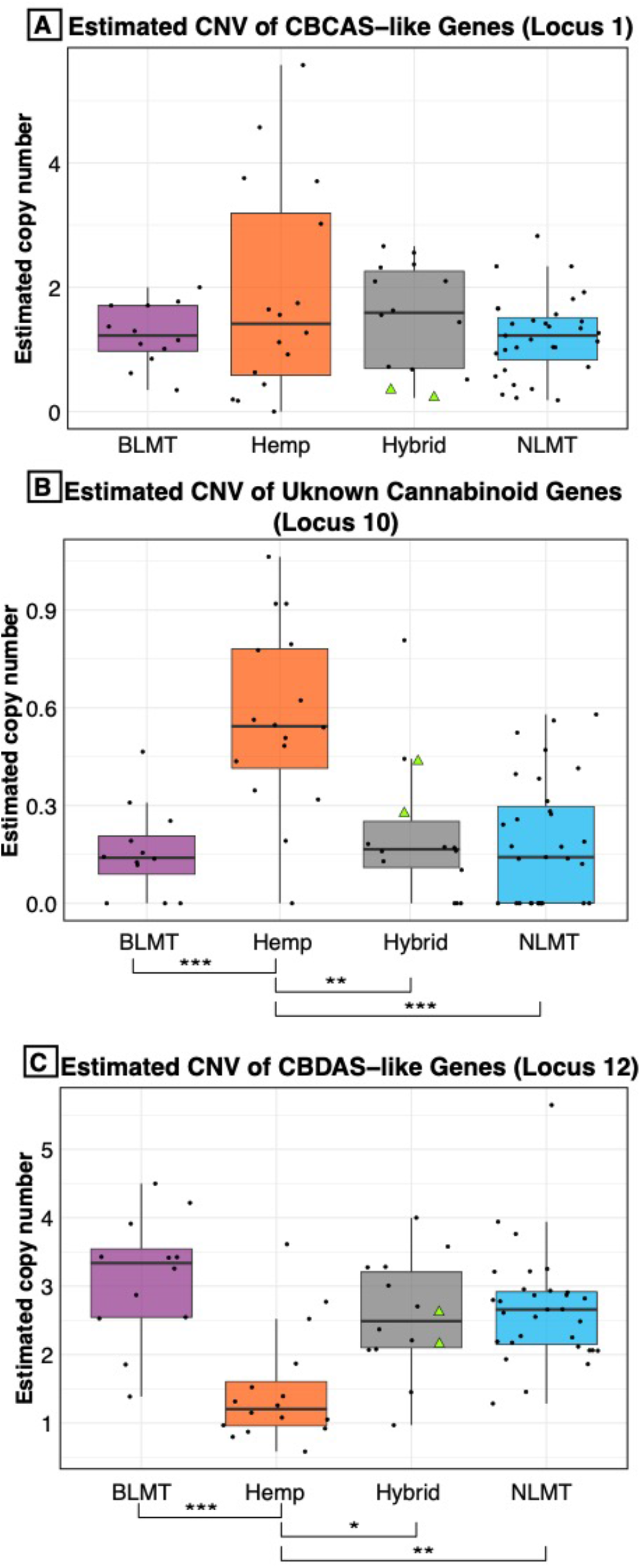
Copy Number Variation in cannabinoid genes. The estimated copy number of the CBCAS-like genes (**A**) is not different between groups despite the hemp lineage having the widest range. Another unknown cannabinoid locus (**B**) shows significant differences between hemp and the other groups at the P< 0.001 level. The hemp lineage also differs significantly with a P< 0.01 from all of the other lineages in the estimated copy number of CBDAS-like genes (**C**). NIDA’s two samples are presented within the hybrid grouping by two green triangles.

The copy number variation for the CBDAS gene was higher, ranging from 1-3 or more copies (BLMT μ=3.24; s.d=1.23; Hemp μ=1.57; s.d=1.04; Hybrid μ=2.59; s.d=1.17 and NLMT μ=2.97; s.d=3.15). The hemp group on average has a lower copy number of these genes, which is significantly different from every other group (Figure 4C). For the CBDAS genes, the first NIDA variety has an estimated copy number of 2.35 and the second one of 2.55. These copy number estimates are close to the mean and median values.

### Maternally inherited genomes

We analyzed both the chloroplast (Figure 5A) and mitochondrial (Figure 5B) haplotype networks. The chloroplast haplotype network (Figure 5A) contains eight haplotypes, with a common haplotype (I) that comprises 58 individuals (79%). Most of the individuals in the haplotypes that diverge from the main haplotype (haplotypes II, V, VI) are hemp types. Both of NIDA’s varieties are part of the main haplotype (I).

**Figure 5.**
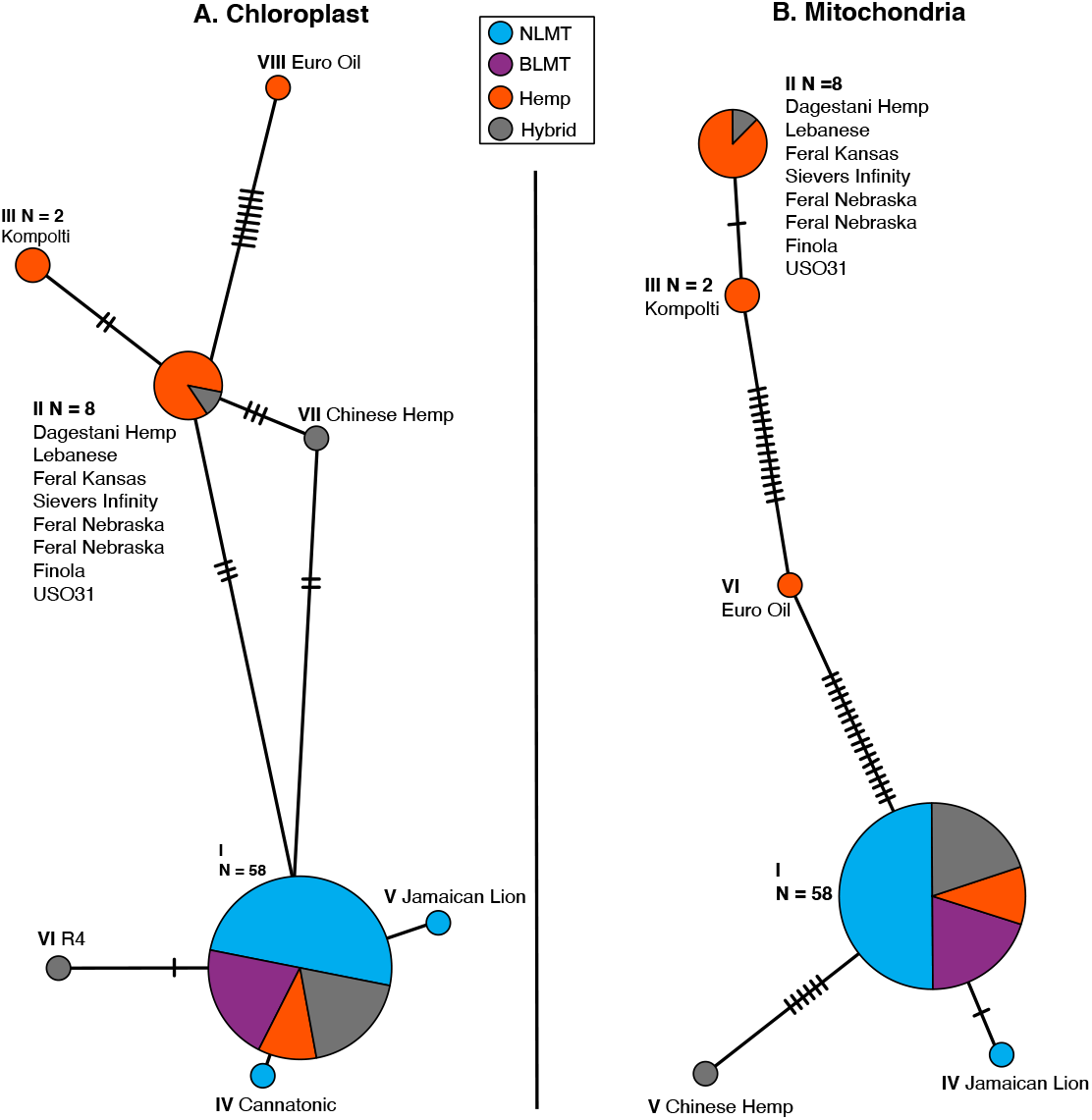
Chloroplast (A) and Mitochondrial (B) haplotype networks. Both haplotype networks are similar with a common haplotype shared by most individuals (79% and 82% for the chloroplast and mitochondria respectively) and smaller haplotypes that differ slightly, mostly comprised of hemp individuals.

The mitochondrial haplotype network has a common haplotype with 60 individuals (82%), and five additional haplotypes which are mostly comprised of hemp individuals (Figure 5B). As with the chloroplast, both of NIDA’s varieties are part of the common haplotype group. The haplotype group for each individual for both the chloroplast and mitochondria is given in columns 11 and 12 in table S1.

### Repetitive genomic content

We found that the 71 genomes analyzed had similar repetitive content in their genomes (BLMT μ=62.9%; s.d=2%; Hemp μ=61.2%; s.d=2.6%; Hybrid μ=62.8%; s.d=2% and NLMT μ=62.9%; s.d=3%) with few outliers (Figure 6). The NLMT had the most variation in the repeat fraction ranging from 58.6% to 70%. Both NIDA samples (showed as triangles in Figure 6) had genome repeat fractions of 61.1%. As showed previously, the majority of repetitive content in *Cannabis* is composed of Long Terminal Repeats (LTR) elements (Ty1 copia and Ty3 gypsy) (Supplementary figure 1).

**Figure 6.**
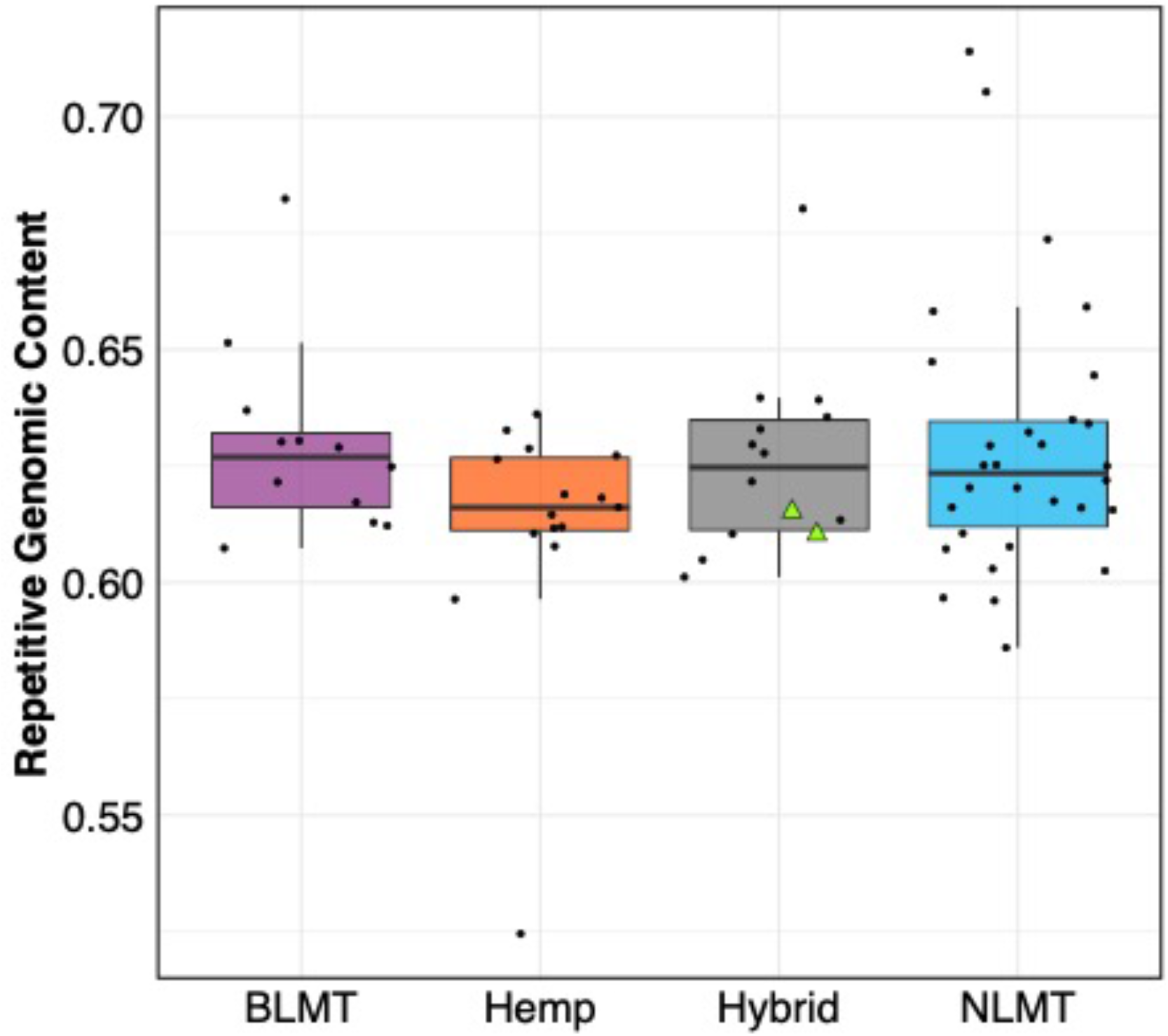
Repetitive Genomic content. The estimated repetitive genomic content by group which does not differ significantly between groups. NIDA’s two varieties are presented within the hybrid grouping by two green triangles.

## Discussion

In this study, we analyzed the genomes of two *Cannabis* samples produced by the sole legal provider of *Cannabis* for research in the U.S., the National Institute on Drug Abuse (NIDA). We compared these two samples to the genomes of 71 commercially available varieties, many of which are medicinally or recreationally consumed by the general public. A previous study has shown that *Cannabis* provided by NIDA lacks diversity and cannabinoid potency compared to commercially available *Cannabis* (Vergara et al. 2017), and microsatellite marker analysis also shows that these differences extend to the genetic level (Schwabe et al. 2019). The results of this study concur with previous studies that NIDA-produced *Cannabis* fundamentally differs from *Cannabis* commonly consumed by the public.

Our whole-genome exploration suggests that the samples from NIDA are very similar to each other, and not divergent to all other varieties in our analysis (Figures 1 and 2), including the varieties commonly used for recreational and medical purposes (Figure 2). Therefore, the samples from NIDA seem to be distantly related to those that are publicly available for consumption.

Even though the two samples supplied by NIDA have high heterozygosity (Figure 3, Table S1), they are comparable to other varieties from the hybrid group and from the NLMT group. The high heterozygosity of both the samples from NIDA could be due to recent outcrossing, and perhaps a recent hybrid origin. However, because we only sampled two individuals, this may not represent the overall heterozygosity of all of the varieties produced for NIDA. Still, as previously stated, research on the chemotypic variation of NIDA’s varieties show the limited cannabinoid diversity (Vergara et al. 2017), supporting the possibility that these two samples are recent hybrids and not bred for their chemotypic profiles including cannabinoids.

The copy number of the cannabinoid genes from the NIDA samples in some cases they fall under the median (Figure 4A), above the median (Figure 4B) or near the median (Figure 4C). However, there are some varieties that have up to 13 copies of some genes (Table S1), in agreement with previous reports (Vergara et al. 2019). Additionally, in the maternally inherited genomes, both NIDA samples have common haplotypes compared to other varieties in the analysis, supporting recent research on the mitochondrial genome diversity in *Cannabis* (Attia et al. 2020). Finally, the repetitive content in the samples from NIDA is comparable to that from other varieties (Figure 6), which is mostly still unknown (Figure S1).

Although the NIDA material used for research does not represent the full range of genetic variation, the results presented suggest that the cannabinoid synthase genes may be similar in many respects to more widely used material. However, the lack of similarity at many other parts of the genome, apparent in the genetic clustering illustrated in Figure 1, may help to explain why the chemistry of NIDA material is so different (Vergara et al. 2017). Differences in cultivation, storage, and processing may also play important roles.

One of the caveats of this investigation is that the Hybrid group is not a lineage of truly related individuals, but a grouping of individuals whose population assignment probability is less than 60% to any of the other groups hence is somewhat arbitrary. Had we chosen a higher Hybrid assignment probability value, there would be fewer individuals in the NLMT, BLMT, or Hemp groupings and more individuals in the Hybrid group. Had we chosen a lower value, there would be fewer individuals in the Hybrid category and more individuals in the other groupings. However, there are individuals with 100% assignment probability to one group, for example, ‘Carmagnola’ has 100% genetic assignment to the Hemp group, ‘Afghan Kush’ has 100% genetic assignment to the BLMT group, and ‘Super Lemon Haze’ has 100% genetic assignment to the NLMT group. If we had chosen a value of 40% instead of 60%, both the NIDA varieties would have grouped with the NLMT group (see Table S2 for the exact assignment probability).

In addition to limiting the research capacity on genetic and chemotypic variation by restricting investigation to only *Cannabis* supplied by NIDA, medical research using this material is also limited and inaccurate. Given that NIDA’s samples do not represent the genomic or phenotypic variation found in *Cannabis* provided by the legal market, consumer experiences may be different from that which is published in the scientific literature. Therefore, medical research is hindered by using varieties that are not representative of what people are actually consuming, making medical research less predictive. The use of NIDA’s *Cannabis* may be one of the reasons why recent reviews have found therapeutic support for three medical conditions (Abrams 2018), while efficacy as an appetite stimulant, as a relaxant, or to treat epilepsy were not supported despite numerous patient reports.

Limiting *Cannabis* types available for study creates an obstacle for scientific discovery. It has been proposed that *Cannabis* may be evolving dioecy from monoecious populations (Divashuk et al. 2014; Razumova et al. 2016; Prentout et al. 2019) and cytonuclear interactions, which could be involved in this transition to dioecy, may be also taking place. To understand processes like these, scientists need access to a diverse and growing variety of *Cannabis* plants which are not available through NIDA. Important discoveries in other plant groups such as transposable elements (McClintock 1950) genes related to pathogen resistance (Leister et al. 1996), or genes related to yield (Sakamoto and Matsuoka 2008) would have not been possible had there been similar restrictions on their research.

This limitation also affects the untapped possibilities of using *Cannabis* to treat a multitude of illnesses, which is backed by a mass of anecdotes. These will continue to be anecdotes until they are studied using rigorous scientific testing methods and scientists are able to provide reliable answers to the community. *Cannabis* is the most widely consumed illicit substance in both in the U.S. and worldwide (Gloss 2014), and therefore it is a matter of public health and safety to provide honest and accurate information. This information is also crucial to policy officials who rely on facts for laws and regulation. In conclusion, scientists must be allowed to use all publicly available forms of *Cannabis* for research purposes in order to maximize scientific, economic, and medicinal benefit to society.

## Author Contributions

D.V. analyzed the single-copy portion of the genome, made figures, wrote the first draft of the manuscript, conceived and lead the project; E.L.H. analyzed the single-copy portion of the genome including STRUCTURE and splits tree graphs, wrote the bioinformatic pipelines; K.G.K. wrote bioinformatic pipelines for the single-copy portion analysis and PCA; R.B.P. analyzed the repetitive content of the genome; A.L.S, M.E.M. acquired DNA samples; and N.C.K. conceived and directed the project. All authors contributed to manuscript preparation.

## Funding

This research was supported by donations to the University of Colorado Foundation gift fund 13401977-Fin8 to NCK and to the Agricultural Genomics Foundation and is part of the joint research agreement between the University of Colorado Boulder and Steep Hill Inc. which made possible the sequencing of the two NIDA genomes.

## Acknowledgments

We thank B. Holmes of Centennial Seeds; D. Liles, C. Casad, A. Ledden and J. Cole of The Farm; MMJ America, Medicinal Genomics, A. Rheingold and M. Rheingold of Headquarters; D. Salama, Nico Escondido, Sunrise Genetics, and B. Sievers for providing DNA samples or sequence information.

## Conflict of Interest

D.V. is the founder and president of the non-profit organization Agricultural Genomics Foundation, and the sole owner of CGRI, LLC. N.C.K. is a board member of the non-profit organization Agricultural Genomics Foundation.

## Supplementary Material

**Supporting Information Figure S1.**
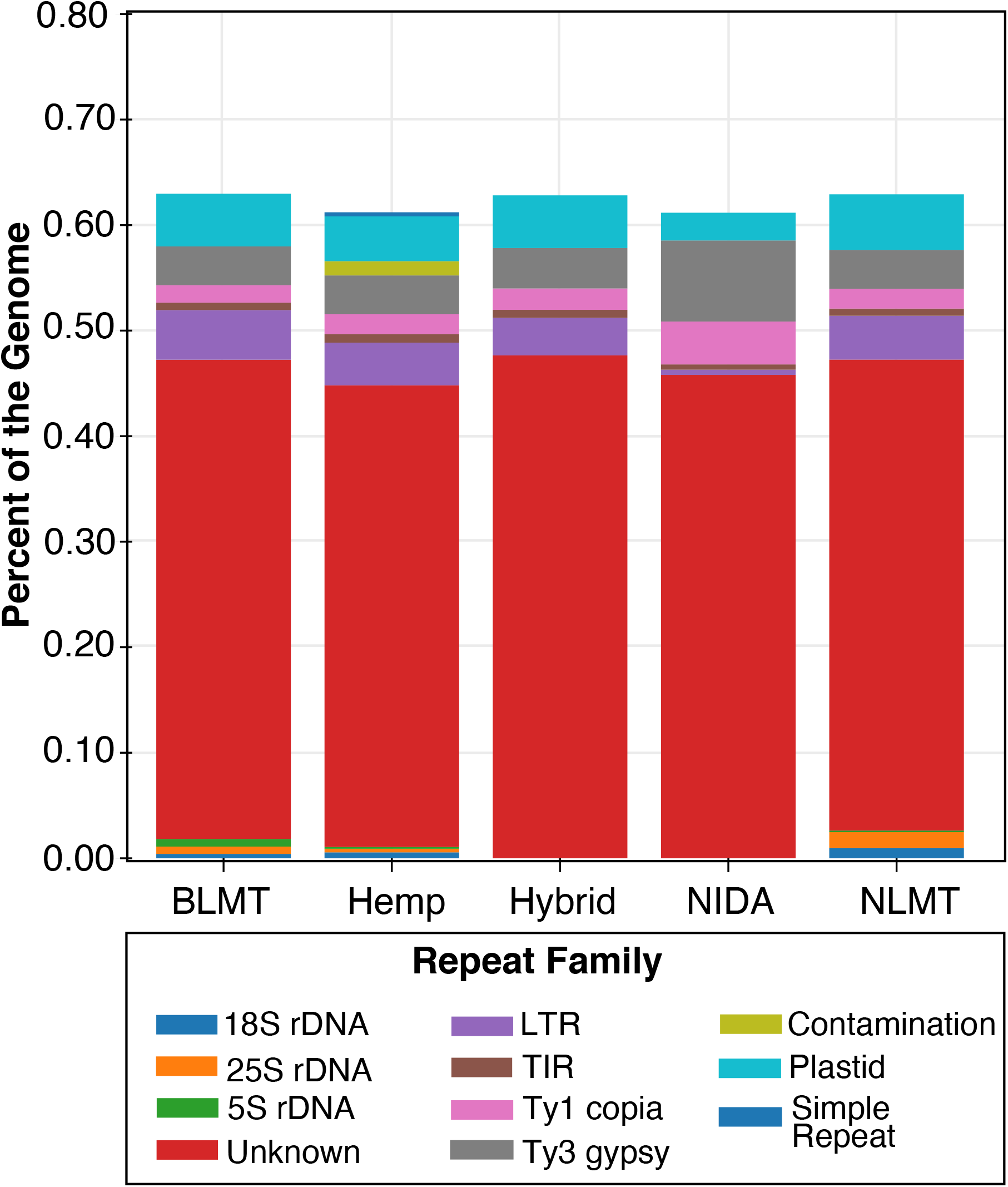
Repetitive content characterization. The graph based clustering algorithm Repeat Explorer characterized the percentage of the genome that belong to the different repeat families. Exact numbers in table S4.

**Supporting Information Table S1. Genetic and genomic information**. Cultivar name (column 1), Sample ID (column 2), classification based on Structure (Column 3), NCBI accession number (column 4), provider (column 5), genome calculations (columns 6-10), haplotype groups (columns 11-12), heterozygosity calculations (columns 13-20), PCA (columns 21-40), cannabinoid loci statistics (columns 41-76).

**Supporting Information Table S2. Population assignment probability**. Sructure’s population assignment probability. Individuals with an assignment probability of <60% to any group were assigned to the ‘hybrid’ grouping.

**Supporting Information Table S3. Cannabinoid BLAST results**. Cannabinoid BLAST results to the Cs10 assembly with more than 80% identity and an alignment length of greater than 1000bp.

**Supporting Information Table S4. Repeat Families**. Different families based on the clustering algorithm used in Repeat Explorer.

